# What’s left in the tank? Identification of non-ascribed aquarium’s coral collections with DNA barcodes as part of an integrated diagnostic approach

**DOI:** 10.1101/2020.07.15.202739

**Authors:** Luigi Colin, Daniel Abed-Navandi, Dalia A. Conde, Jamie Craggs, Rita da Silva, Max Janse, Björn Källström, Alexander Pearce-Kelly, Chris yesson

## Abstract

The unprecedented threats to coral reef ecosystems from global climate change (GCC) require an urgent response from the aquarium community, which is becoming an increasingly vital coral conservation resource. Unfortunately, many hermatypic corals in aquaria are not identified to species level, which hinders assessment of their conservation significance. Traditional methods of species identification using morphology can be challenging, especially to non-taxonomists. DNA barcoding is an option for species identification of Scleractinian corals, especially when used in concert with morphology-based assessment. This study uses DNA barcodes to try to identify aquarium specimens of the diverse reef-forming genus *Acropora* from 127 samples. We identified to our best current knowledge, to species name 44% of the analysed samples and provided provisional identification for 80% of them (101/127, in the form of a list of species names with associate confidence values). We highlighted a sampling bias in public nucleotide sequences repertories (e.g.: GenBank) towards more charismatic and more studied species, even inside a well-studied genus like *Acropora*. In addition, we showed a potential “single observer” effect with over a quarter of the reference sequences used for these identifications coming from the same study. We propose the use of barcoding and query matching as an additional tool for taxonomic experts and general aquarists, as an additional tool to increase their chances of making high confidence species-level identifications. We produce a standardised and easily repeatable methodology to increase the capacity of aquariums and other facilities to assess non-ascribed species, emphasising the value of integrating this approach with morphological identification optimising usage of authoritative identification guides and expert opinion.

## Introduction

It is widely acknowledged that coral reef ecosystems are facing unprecedented threats from global climate change (GCC) e.g.(Pratchett et al. 2013; Hoegh-Guldberg et al. 2017; Cowburn et al. 2018; Hughes et al. 2018; Masson-Delmotte et al. 2018; Brondizio et al. 2019; Sheppard et al. 2020). Warm water reef-building corals create the most biodiverse ecosystems on the planet and have been estimated to support 830’000 species of multi-cellular plants and animals worldwide (Bellwood and Hughes 2001; Mora et al. 2008, 2011; Fisher et al. 2015), providing a variety of habitats for fish, invertebrates and other taxa in shallow tropical seas (Bellwood and Hughes 2001). *Acropora* corals play a key functional ecosystem role as they are the major reef builders in the majority of warm water reefs ecosystems (Fukami et al. 2000).

### Corals in aquaria

*Ex situ* conservation can play an important role in ensuring the survival of many species (Turley 1999; Hutchins 2003; Raven 2004; Blanco et al. 2009; Fa et al. 2011; Leus et al. 2011; Lacy 2013; Skibins and Powell 2013). Given the severity of the species and ecosystem level climate change threats to corals, and the alarming rates of recent losses (Masson-Delmotte et al. 2018), coral collection can potentially play a valuable role in future conservation efforts. Aquaria institutions therefore have a vitally important conservation remit to support coral restoration strategies allowing for both sexual and asexual coral recruitment, biobanking and a wide range of research, education and thus contribution towards *in situ* conservation activities. Early attempts in the 1980s at keeping corals in aquaria faced difficulties (Borneman 2008; Brunner 2012) it was until 1980 that *Acropora* was the first genus to be successfully cultured by Stüber (Borneman 2008). From the 1990s onwards coral husbandry boomed, reaching a level where virtually all families of zooxanthellate corals were not only being maintained for many years, but were being propagated and traded between private and public aquaria (Borneman 2008). Specimens conserved in Aquaria thus, became a highly valuable resource for restoration strategies (Rinkevich 1995).

### Identification of corals

Coral taxonomy is a traditionally difficult discipline, subject to all the challenges of modern taxonomy (Godfray 2002). The genus *Acropora* is among the most diverse and geographically widespread reef building corals (Wallace 1999). Research on *Acropora* suggests that species limits may be sometimes narrower, sometimes broader, than generally perceived (Wallace and Willis 1994). Somewhat blurry species boundaries are confounded by hybridisation (Willis 1997; Hatta et al. 1999; Van Oppen et al. 2002; Vollmer and Palumbi 2002; Wolstenholme 2004; Willis et al. 2006; Palumbi et al. 2012; Wei et al. 2012; Isomura et al. 2013), often worsened by the possibility of synchronised spawning events (Harrison et al. 1984; Babcock et al. 1986; Márquez et al. 2002; Van Oppen et al. 2002). Currently the number of officially recognised *Acropora* species are between 135 (WORMS (Horton et al. 2020)) and 163 (Corals of the World (Veron et al. 2020)), reaching 186 if we include the *“taxon inquirendum”* on WORMS (Horton et al. 2020). There have been different attempts at grouping these species within the genus *Acropora* (Wallace 1999). These attempts lead to the creation of the concept of species group (aka syngameon). The concept of species groups, first defined for convenience of identification and without implying taxonomic relation, has been revised to reflect the evolution and phylogeny of the genus (Wallace 1999; Wallace et al. 2012). The use of genetic tools for species identification has provided another line of evidence for species delimitation, further adding to the blurred boundaries between species groups, species, and sub-species.

Traditionally, identification of coral has relied on morphological skeletal features, with the great majority of taxa originally described following typological species concepts defined around a century ago (Best et al. 1984; Todd 2008). The relationship between morphological variation within-species and the environment has been a source of contention since the late 1800s (Todd 2008). Whether this variability is due to different underlying genotypes or plastic phenotypes is unclear. Evidence supports both scenarios (difference in phenotype e.g.: (Willis 1985; Ayre and Willis 1988; Todd et al. 2004) Plastic phenotypes e.g.: (Foster 1979; Miller 1994; Bruno and Edmunds 1997; Muko et al. 2000; Todd et al. 2004)) and these scenarios are not necessary mutually exclusive but probable to operate simultaneously (Foster 1979; Amaral 1994; Todd 2008). This has particular relevance in the face of continuous GCC, that is creating an increasingly unpredictable wild reef environment.

Notwithstanding the taxonomic difficulties, the identification of coral species can be challenging even in optimal ‘wild’ conditions. However, identification requirements are far more than just for well documented field collections, but also cover aquaria collections, that in some cases are held in conditions dramatically different from the wild. Furthermore, customs officials rely in proper taxonomic identification even at the genera level for threatened taxa by the international wildlife trade (AC25 Doc. 23 CITES 2011), although currently *Acropora* are covered by the blanket listing of all Scleractinia under CITES appendix II. A status of these identification difficulties in aquaria can be demonstrated by an analysis of Species360 Zoological Information Management Software (ZIMS), adopted by more than 95 aquariums in 24 countries. Assuming correct identification ZIMS indicates c. 42.9% of corals of the orders Scleractinia, Alcyonacea, Helioporacea, Antipatharia, Corallimorpharia, Pennatulacea, and Zoanthidea in aquaria are identified to genus or higher taxonomic level (ZIMS list of species holdings – 4th June 2020). Most of the records correspond to the order Scleractinia, which is also the order with more species represented in aquariums. Corallimorpharia is, however, the order with the highest number of reported individuals in aquariums members of the Species360 network (Fig. 1). In addition to this the non-species associated genera (e.g.: *Acropora* sp.) is usually recorded once in the ZIMS database and most other aquarium inventory systems, which possibly significantly underestimates the actual number of non-identified specimens. Moreover, there are currently no confidence values ascribed to any taxonomic level a collection allocates to a specimen which means that there is likely to mean that a significant number of species ascribed specimens are actually at the lower end of the confidence spectrum and would benefit from being reassessed.

**Fig. 1:**
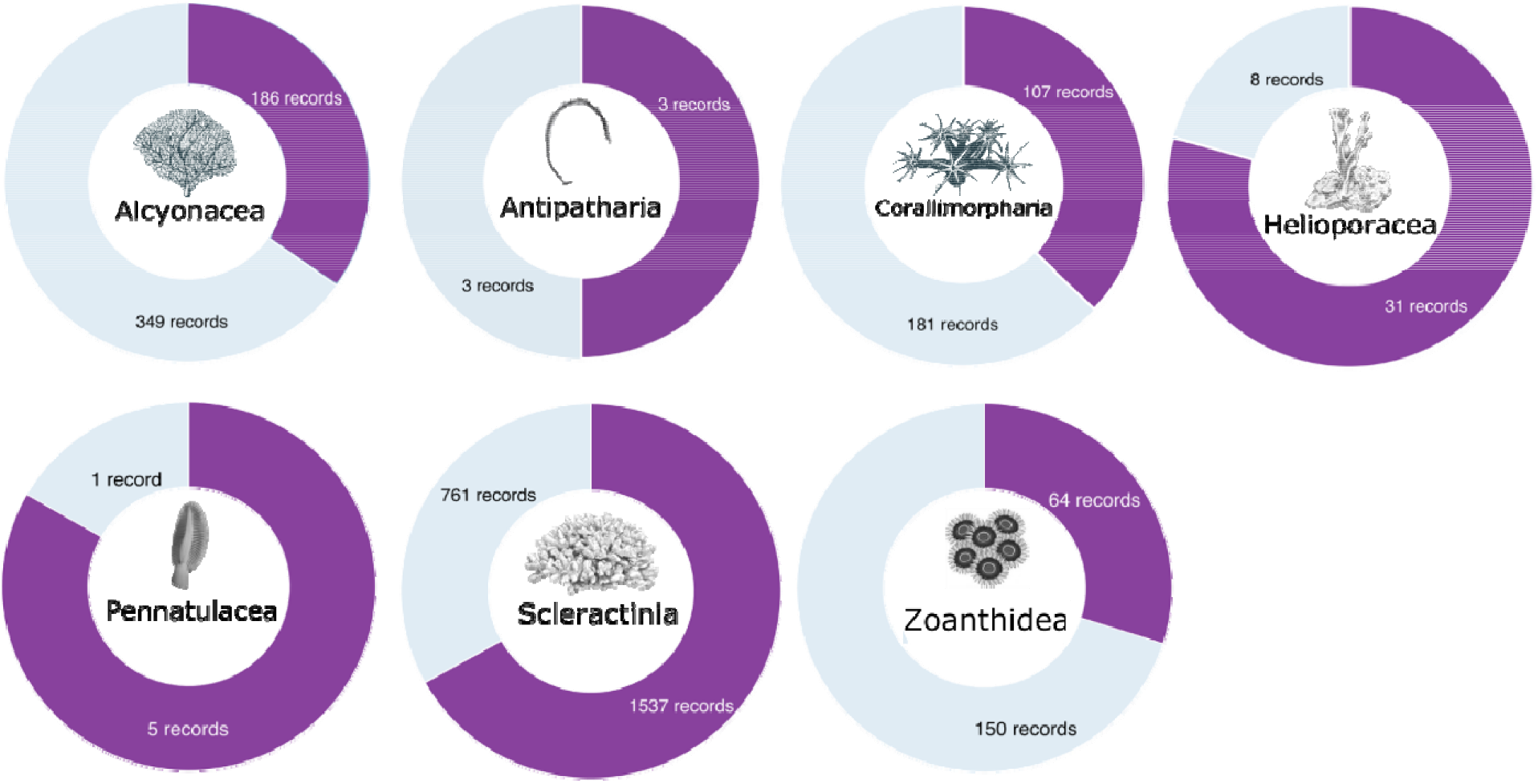
Number of records from member institutions of the Species360 network at genus or higher level in light blue (i.e. order, family and genus) and at species or lower level in purple (i.e. species and subspecies). This information is based on the rank field from the holdings information reported by each institution through the ZIMS software. Information downloaded from ZIMS (Species360, 2020) on 4th June 2020.

In many aquariums’, corals come from three sources: fragments from other aquaria, via ornamental trade, or from confiscated and seized shipments (which aquariums are usually asked to accommodate). All three sources present identification challenges as often exact knowledge from their wild origin is absent. This specially a challenge for confiscated specimens since they have none or relatively poor source records, sample integrity, and any other additional background information that might be of assistance for the identification process. This, together with the difficulties caused by environmental phenotypic plasticity makes coral identification in aquariums a particularly challenging task. Data from the Convention on the International Trade of Endangered Species of Flora and Fauna (CITES) shows a total of 4557 instances of coral confiscation from 1982 to 2018 worldwide, for a total of 27027 individual items and 5461 kg per year (CITES data - https://trade.cites.org02/07/2020). Of these, less than 20% are reported at species level, meaning that for most cases we know only the genus, family or even order of the specimens confiscated. Given these challenges CITES currently requires identification only to the order level.

### DNA Barcoding

An alternative or complementary diagnostic option to morphological identification is genetic analysis. DNA Barcodes have been proposed by Hebert et al. (Hebert et al. 2003) as a way to circum-vent the “limitations inherent in morphology-based identification systems”. The core concept of DNA barcoding is the use of a standard sequence corresponding to a single homologous region that can be amplified with universal primers that is sufficiently variable to distinguish between species – usually represented by the mitochondrially encoded cytochrome c oxidase I (MT-CO1 or CO1 or COX1) (Hebert et al. 2003).

Beside the inherent limitations (Shearer et al. 2002; Ferguson 2002; Moritz and Cicero 2004; DeSalle et al. 2005; Meyer and Paulay 2005; Prendini 2005; Cognato 2006; Hickerson et al. 2006; Meier et al. 2006, 2008; Little and Stevenson 2007), DNA barcoding is especially attractive as an option for species identification of scleractinian corals, as this method bypasses the problems caused by phenotypic plasticity or life stage (larva, juvenile or adult). Unfortunately, the proposed ‘universal’ Barcoding region CO1 is at least 10-20 times slower in Anthozoa than the standard vertebrate mtDNA mutation rate, and 2-5 times slower than the average Anthozoan nuclear sequence(Van Oppen et al. 1999; Shearer et al. 2002; Hellberg 2006; Huang et al. 2008; Chen et al. 2009; Ortman et al. 2010) with zero interspecific divergence for many species (Shearer and Coffroth 2008). This has led to the use of a number of different barcoding regions for corals (Shearer et al. 2002).

The diverse, reef forming Acropora group are widely grown in aquaria and have been a target for barcoding work. Potential barcodes were reviewed by Shearer et al. (Shearer et al. 2002) examining both nuclear and mitochondrial regions. Substitution rates of mitochondrial genes in *Acropora* tend to be lower relative to the nuclear counterparts mentioned by Shearer (i.e. Pax-C intron; internal transcribed spacer 1 and 2 (ITS) (Shearer et al. 2002). Although, Van Oppen et al. (van Oppen et al. 1999) and Vollmer et al. (Vollmer 2002) have highlighted the mitochondrial putative control region (mtCR) is more variable than the other mitochondrial regions. Subsequently Shearer et al. (Shearer et al. 2002) compared the genetic divergence of mitochondrial and nuclear regions in Scleractinia, highlighting how the mtCR region has a similar range of divergence to nuclear coding regions like the Pax-C intron, but is much slower than the hyper-variable ITS1-2. Notwithstanding all the above mentioned work, *Acropora’s* molecular taxonomy remains to be resolved, and to do so it would require extensive taxonomic work and genome-wide markers (Cowman et al. 2020).

This study seeks to build on these studies by investigating the utility of DNA barcoding as a method for provisional species identification of *Acropora* collections in aquaria, as an aid to morphological taxonomy rather than an alternative, and as part of an integrated diagnostic approach.

## Material and Methods

### Study Area and Collection

As part of the “idCoral” project we collected 224 samples across seven institutions (Aquarium de Paris (France), Chester’s Zoo (UK), Haus des Meeres-Vienna (Austria), Horniman Museum and Gardens (UK), Royal Burgers’ Zoo (Netherlands), Tierpark Hagenbeck Aquarium (Germany), ZSL London Zoo (UK)). Branch tip fragments of 2 cm were collected and stored in single ethanol (+95%) vials labelled with a unique identifier. During the sample collection the colonies were tagged and photographed with collection dates and tank locations were recorded. The photographs and tags were also recorded in the “idCoral” database (database - idcoral.org (2020)). After collection samples were stored at −20°C until extraction, with an initial change of ethanol after 1-2 days aimed at reducing degradation. We analysed 127 *Acropora* samples from these collections before COVID-19 restrictions stopped further progress on this project.

### DNA Extraction, Amplification and Sequencing

The DNA extractions were undertaken using the DNeasy PowerSoil Pro Kit from Qiagen^©^ with a modified version of the manufacturer’s protocol. An incubation step with proteinase K was added and the physical breaking of the cell structure with a chemical dissolution was removed from the protocol as it resulted in excessive shearing of the DNA (Online Resource 1). Extracted DNA was then preserved at −20°C, and the DNA concentration and quality tested with both Nanodrop assay and gel electrophoresis. The DNA was diluted to standardise the concentration to between 5 to 20 μg/ml.

Two genes, PaxC and mtCR were selected as barcodes based on an assessment of their variability and availability as reference sequences (Van Oppen et al. 2001; Shearer et al. 2002). The ITS1-2 region was excluded due to the lower number of taxa available as reference sequences. PCR amplification was conducted with Q5® High-Fidelity DNA Polymerase mixed following manufacturer’s indication. 3μl of DNA template and 0.5 μl of Bovine Serum Albumin (BSA)1μg were added, reaching a final volume of 25μl with RNA free water. For the PaxC gene the primer used were PaxC FP1 5’-TCC AGA GCA TTA GAG ATG CTG G-3’ and PaxC RP1 5’-GGC GAT TTG AGA ACC AAA CCT GTA-3’ (Van Oppen et al. 2001) with a protocol of 98°C for 30s, followed by 31 cycles at 98°C for 10 s, 62°C for 30 s and 72°C for 60 s, ending with a final phase of 72°C for 2min. For the mtCR the primer used were RNS2: 5’-CAG AGT AAG TCG TAA CAT AG-3’ and GR: 5’-AAT TCC GGT GTG TGT TCT CT-3’ (Suzuki et al. 2008) with a protocol of 98°C for 30s, followed by 40 cycles at 98°C for 10 s, 62°C for 15 s and 72°C for 60 s, ending with a final phase of 72°C for 5min.

Successful PCR amplifications (assessed by gel electrophoresis) were sent to Eurofins Genomics for PCR cleaning and custom Sanger sequencing on both the forward and the reverse primers using cycle sequencing technology (dideoxy chain termination / cycle sequencing) on ABI 3730XL sequencing machines. The returned sequences were manually trimmed to remove poor quality sections and assembled using Geneious Prime^®^ (2019.1.2.).

### Identification

#### Taxonomy

Both the barcode gap and specimen identifications were initially performed at the species level, but were also performed at the level of species group, based on the revision of Acropora by Wallace et al. (Wallace et al. 2012), to provide an additional layer of information in the final integrated approach.

#### Barcoding Gap

Reference sequences for the two targeted regions were collated. FASTA files were created with the records matching the following query on GenBank: mtCR=““Acroporidae”[Organism] AND mitochondrial “complete genome” OR “control region” OR “putative control region” OR mitochondrial control region”; PaxC=““Acroporidae”[Organism] AND PaxC[gene] OR pax-c[gene]”. Two multiple alignments were performed, one for each gene, using the Geneious^®^ algorithm (progressive pairwise aligner) and trimmed. For both the produced references alignments distance matrices were calculated using the function dist.dna (R package apex (Jombart et al. 2017)) and with different models (Raw; JC69; K80; K81; F84; BH87; paralin; indel Y; indelblock). Successively the frequency distribution of intra/inter-specific distances was plotted. The quantile function (R package “stats”) was used to define the 99; 50; 1% quantile confidence interval used in the blast match analysis. Note: the two alignments were constructed and used only for the specific purposes of examining the barcoding gap for the targeted region and determining potential confidence interval for the blast match analysis.

#### Phylogenetic tree

Separate unrooted Maximum-likelihood (ML) phylogenetic trees were created, one for each targeted region. The references obtained from the previously mentioned query were cleaned for duplicated sequences and subsequently aligned. The two phylogenetic analyses were performed using the program IQ-TREE, with default settings and automatic model selection (Minh et al. 2013; Nguyen et al. 2015) with analysis conducted on the usegalaxy.eu public server (Afgan et al. 2018).

#### Blast match

Query based specimen identification with blast match (Benson 2000) was determined using a custom Blastn database. All records from the query ““Acroporidae”[Organism]” on GenBank were downloaded and included in the local database built with the packages rBLAST (Hahsler and Nagar 2019) and rentrez (Winter 2017). A local Blast match (Camacho et al. 2009) was ultimately performed. To filter for species match with identical scores a decision tree (Fig. 2) was designed relying on Bit score, Percentage identity, and number of mismatches. Matches shorter than 2/3 of the original sample sequence query were removed. The decision tree prioritised 100% pairwise identity matches where possible, then to the highest bit score, then lowest number of base mismatches. 100% pairwise identity matches to a species group rather than a single species name were prioritised over single species name matches with lower pairwise identity with higher bit score, both possible IDs were retained. We confronted the match from the two targeted regions to obtain a single most probable ID. To account for conflicting matches between the two region we relied on the 99; 50; 1% quantile distribution of the intraspecific distances obtained from the barcoding gap analysis. Using pairwise identity threshold as confidence intervals to discern which region offers the better match for the ID (R-scripts as Online Resource 3 and 4).

**Fig. 2:**
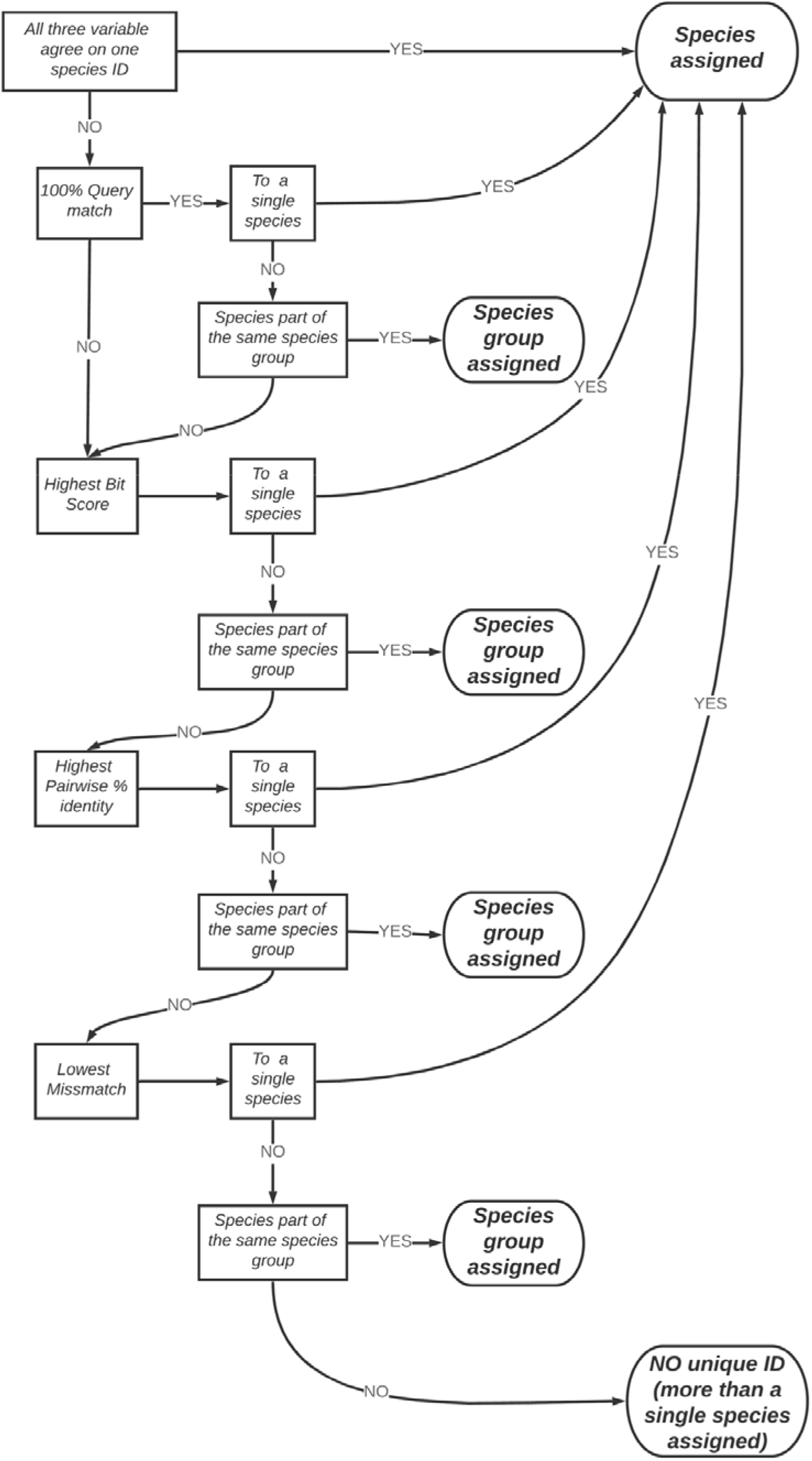
Decision tree for conflict solving based on Pairwise percentage identity, Bit score and number of mismatches.

## Results

From the 127 treated samples all were successfully extracted and, 107 mtCR and 100 PaxC had a successful DNA amplification, respectively 84% and 79% with 76 samples overlap between the two regions (Table 1). 348 reads were successfully sequenced, 84 mtCR sample and 90 PaxC. The median sequence lengths were 734 post assembly for the mtCR region and 649 for the PaxC region (Table 1).

**Table 1:**
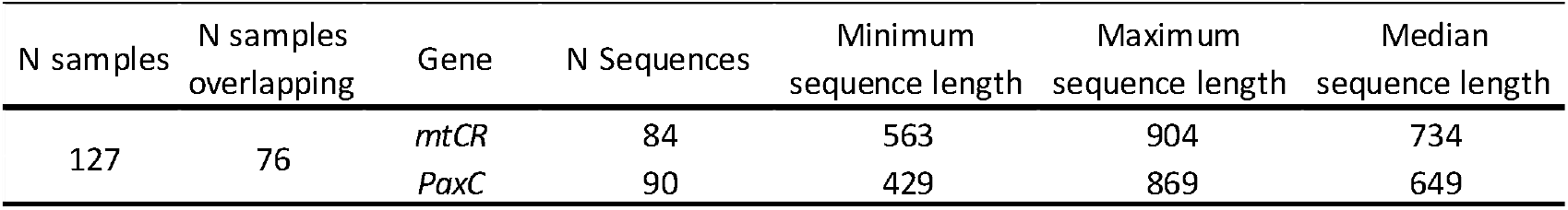
Success rate across genes

| N samples | N samples overlapping | Gene | N Sequences | Minimum sequence length | Maximum sequence length | Median sequence length |
| --- | --- | --- | --- | --- | --- | --- |
| 127 | 76 | *mtCR* | 84 | 563 | 904 | 734 |
|  |  | *PaxC* | 90 | 429 | 869 | 649 |

### Barcoding

Our GenBank query produced a total of 1348 mtCR sequences and 427 PaxC sequences. The mtCR references query, once all sequence duplicates with the same species name were removed, produced a 579-sequence alignment of length 914bp. The PaxC references query, after similar cleaning produced a 352-sequence alignment of length 770 bp.

The mtCR alignment of 579 sequences resulted in a matrix of 335 241 pairwise genetic distances, grouped by 57 species names. 6.4% of these (21 471) are intraspecific distances, while 93.6% (313 770) are interspecific distances (Fig. 3). The PaxC alignment of 352 sequences resulted in a matrix of 123 904 genetic distances, grouped by 55 species names. 3% of these (3 726) are intraspecific distances, while 97% (120 178) are interspecific distances (Fig. 3). Between the two groups there is an overlap of 49 species names. Table 3 and 4 with the sources of the sequences used in these alignments.

**Fig. 3:**
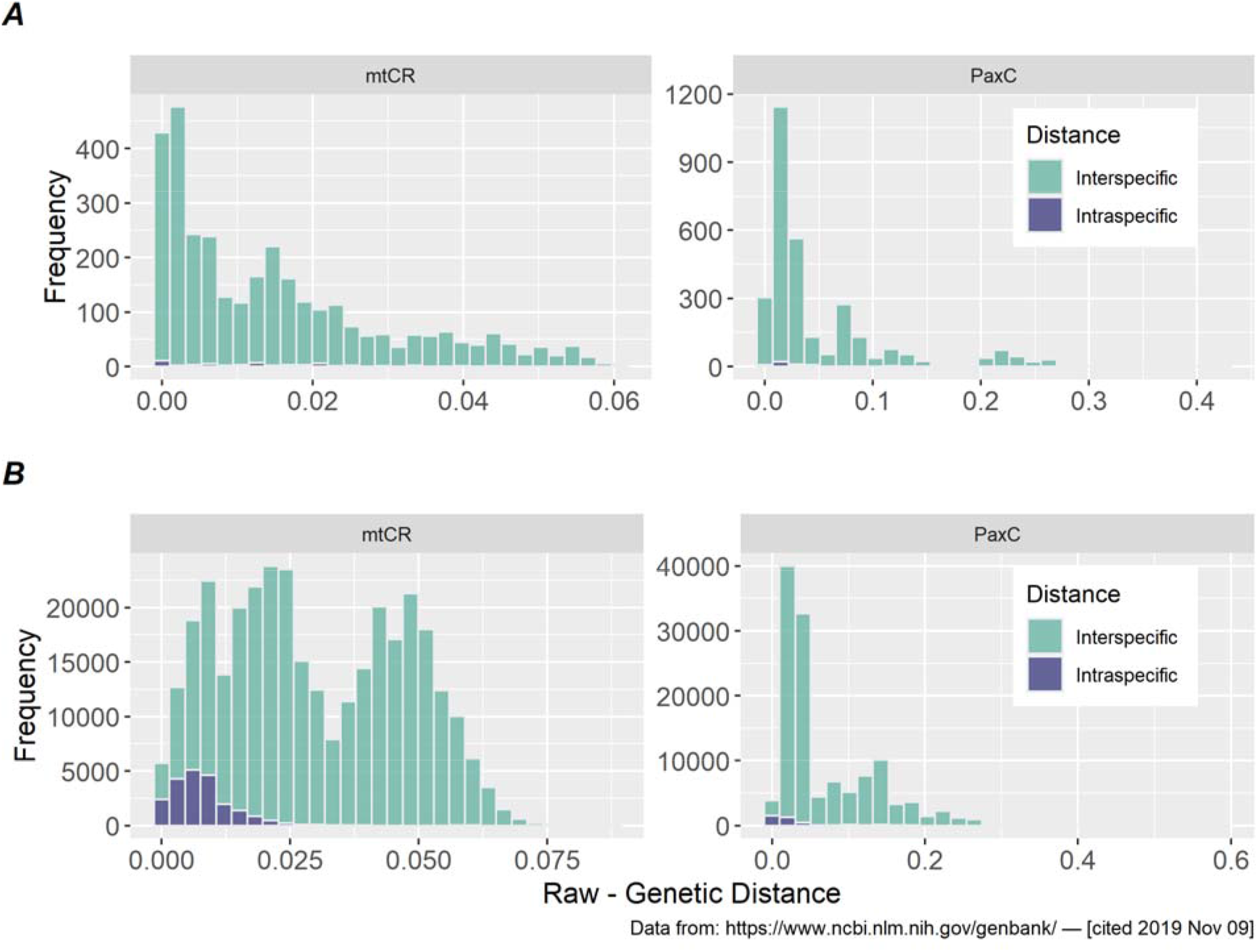
Barcoding gap (raw distance matrix frequency distribution) – mtCR max distance = 0.0877 minimum distance = 0; PaxC max distance = 0.5895 minimum distance = 0. (Bin size=0.002) A) Minimum interspecific distance and maximum intraspecific distance B) all distances

Fig. 3 highlights in blue the intraspecific variation (sequences with the same name on GenBank) and in the light green the interspecific distances (sequences with different names on GenBank). In both Fig. 3A) and 3B), and both barcoding regions, we see no discernable barcoding gap.

The quantile within species distribution produce the following distance threshold for our confidence interval: mtCR 0.027 (99%); 0.0068 (50%); 0 (1%) and PaxC 0.17 (99%); 0.014 (50%); 0 (1%).

In Fig. 4, for the mtCR and the PaxC respectively, we can see the above-mentioned uneven distribution of the GenBank records towards specific species and species groups. With example like species group *humilis* showing 4 times the frequency of less represented species group like *loripes*.

**Fig. 4:**
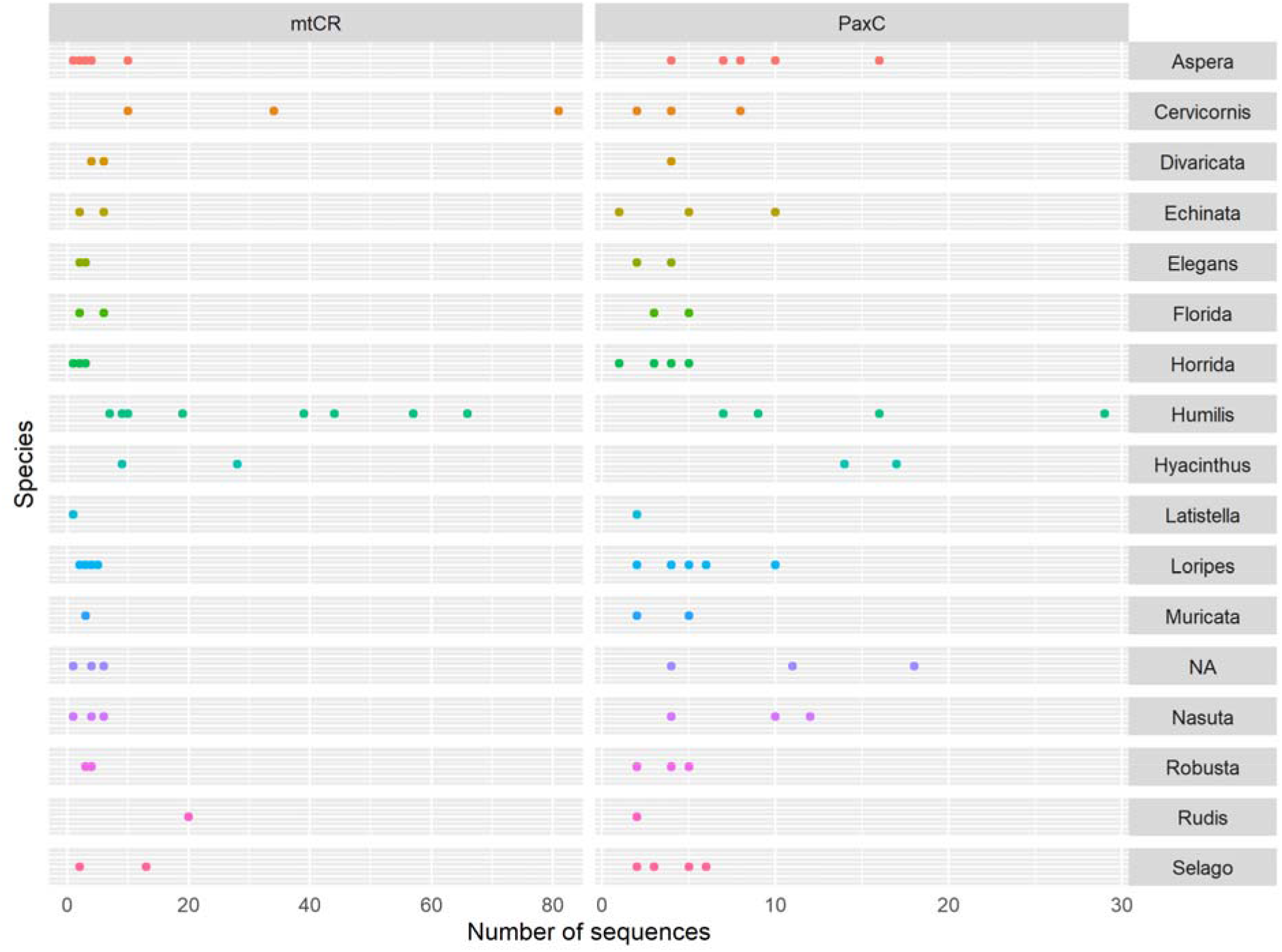
Number of mtCR and PaxC references sequences with the same species name by species group

### Phylogeny

Both the mtCR (Online Resource 5) and the PaxC (Online Resource 6) phylogenies show clear sign of polyphyly. Looking at our best matches (i.e.: 100% match to a reference), most of our sample sequences match to single species – 63% of mtCR 100% matches and all of the PaxC 100% matches. Focusing on just these sequences with 100% matches to a reference, we see that 0/19 mtCR sequences and 1/4 PaxC match to single-species clades (a monophyletic group composed of more than one sample of a single taxon name). However, some of these match to species that a polyphyletic within the tree (All of 100% mtCR matches tie to polyphyletic taxa and 75% for PaxC).

### Blast match

The local database for blast matching contained 510826 sequences. In Table 2 we see that 23 of our 76 samples (with both regions sequenced) were uniquely identified to a single species, while 13 samples matched to conflicting names. 20 of our samples had a unique species name match for the PaxC region but mtCR region matched to 2 or more names with all but 6 of these including the matching PaxC name. Very few samples matched to multiple species for both regions (4).

**Table 2:**
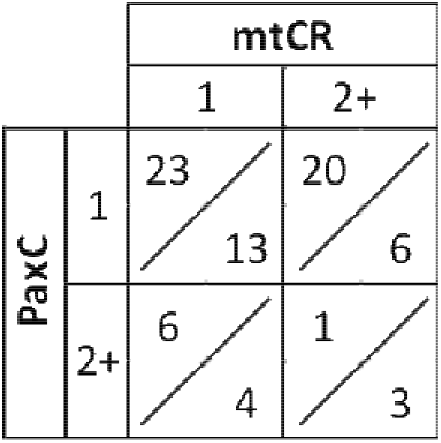
Number of sample matching to a single or more (2+) species name on GenBank. Above the diagonal line match that agree between the two region. Below the diagonal line matches that disagree between the two regions

We have 47 % of samples matching to a single species name with both regions, 5% with the same identical match. While we have 42% of the matches to a single species name that are in conflict, 25% of these conflicts are resolvable (i.e.: one of the two region with a match better fitting than the other one, based on the defined confidence thresholds of each region, rather than directly on the pairwise identity). Similar results for the other sample matching for two or more in at least one of the two regions. Out of the 76 samples, in 15 cases we observe the second region adding information to the first ones (Table 5 as Online Resource 2). Mainly from a mtCR multi name ID to a single PaxC name (i.e.: sample P269 has 7 mtCR identical matches to 7 different species names, and the PaxC matches at a lower rate to only one of them). In addition, the second region functions as backup option whenever one of the two falls too short (5/76 sample have too low-quality match to be trusted independently).

The complete list of our current best species name match, with species group information and detailed genetic info is provided as Online Resource (Table 5 – Online Resource 2).

## Discussion

### Barcoding gap

Using DNA barcodes to identify unknown samples is only achievable if a well-studied, well-sampled reference sequence database is available (Meyer and Paulay 2005), and even though databases such as GenBank (which comprises the DNA DataBank of Japan (DDBJ), the European Nucleotide Archive (ENA), and GenBank at NCBI - http://www.ncbi.nlm.nih.gov/genbank) and BOLD (Barcode Of Life Data System - http://www.boldsystems.org) contain an ever increasing number of species-level identified sequences (i.e.: 216 531 829 total sequences on GenBank as April 2020), these databases are far from complete. The open nature of GenBank is destined to produce a bias towards species with a higher research interest (i.e.: Human sequences constitute 56% of the total sequences on GenBank (Benson 2000)). The genus focus of our study, *Acropora*, is one of the more commonly studied and widespread hard corals in the ocean (Wallace 1999; Fukami et al. 2000) making it one of the most covered coral genera in GenBank. The reality is that even for a widely studied group such as *Acropora* we can observe a disparity in the number of sequences available for each species name (Fig. 4). The best example of this is the delta between two species group: group *humilis* being the most sampled (251 mtCR and 61 PaxC references sequences from our query) and group *robusta* one of the least sampled (11 mtCR and 13 PaxC references sequences from our query). Note that both species group are formed by of the same number of species (8 species each (Wallace et al. 2012). However, while counting the number of sequences available for a group may be an indication of the level of research that group receives, it does not mean that groups with the most sequences have the best reference collection for a DNA barcoding project. The most important quality for a reference sequence collection is it’s representation of the underlying species diversity, ideally covering a wide geographical and morphological range (Vellend and Geber 2005; Rozenfeld et al. 2007; Ma et al. 2017), all-the-while using a marker that shows sufficient variability to permit robust identifications. Sampling bias is a problem for all specimen collections and this must always be kept in mind when interpreting results (Leray et al. 2019).

From our analysis we could not identify a clear barcoding gap in both our targeted regions (Fig. 3), and furthermore we can observe a total overlap of the intra-specific distances by the inter-specific distances. For both our targeted regions, even if more evident in the mtCR (Fig. 3A), we see that the inter-specific distances follow a distribution reminiscent of a more common and more expected barcoding distribution (Fig. 3B, mtCR). We performed the same barcoding gap analysis at the higher taxonomic level using the species group information without obtaining a noticeable difference. With the lack of a clear Barcoding gap, and accounting for the extensive criticism present in the literature: For the use of distances in taxonomy and systematics (Ferguson 2002; Lee 2004; Moritz and Cicero 2004; DeSalle et al. 2005; Prendini 2005; Knapp et al. 2005; Meyer and Paulay 2005; Cognato 2006; Hickerson et al. 2006; Little and Stevenson 2007; Meier et al. 2008); for the drawbacks of using GenBank data (Meier et al. 2006); for the evidence of different rates of evolution in Scleractinia (Van Oppen et al. 1999; Shearer et al. 2002; Kitahara et al. 2010). We cannot establish a simple distance base threshold to use for our identification purposes, leading us to depend more on the blast match.

Another potential issue with the use of these methods of identification is the source of the reference sequences. With databases like GenBank, with little or no review on the uploaded information, the question is the confidence that the initial identification of the now available reference sequences is indeed correct (James Harris 2003). In our specific case after the processing we performed for the mtCR reference alignment, out of the total 579 included 42% (242 sequences) come from a single doctoral thesis project, while 5.4% (33 sequences) are from unpublished papers (Table 3). For the PaxC 16% (58 sequences) are from unpublished papers, and another 39% (138 sequences) from a single paper (Richards et al. 2008) (Table 4). Highlighting clearly the need for more study on the subject to increase the number of potential sources and alleviate the possible bias of a “single observer”.

**Table 3:**
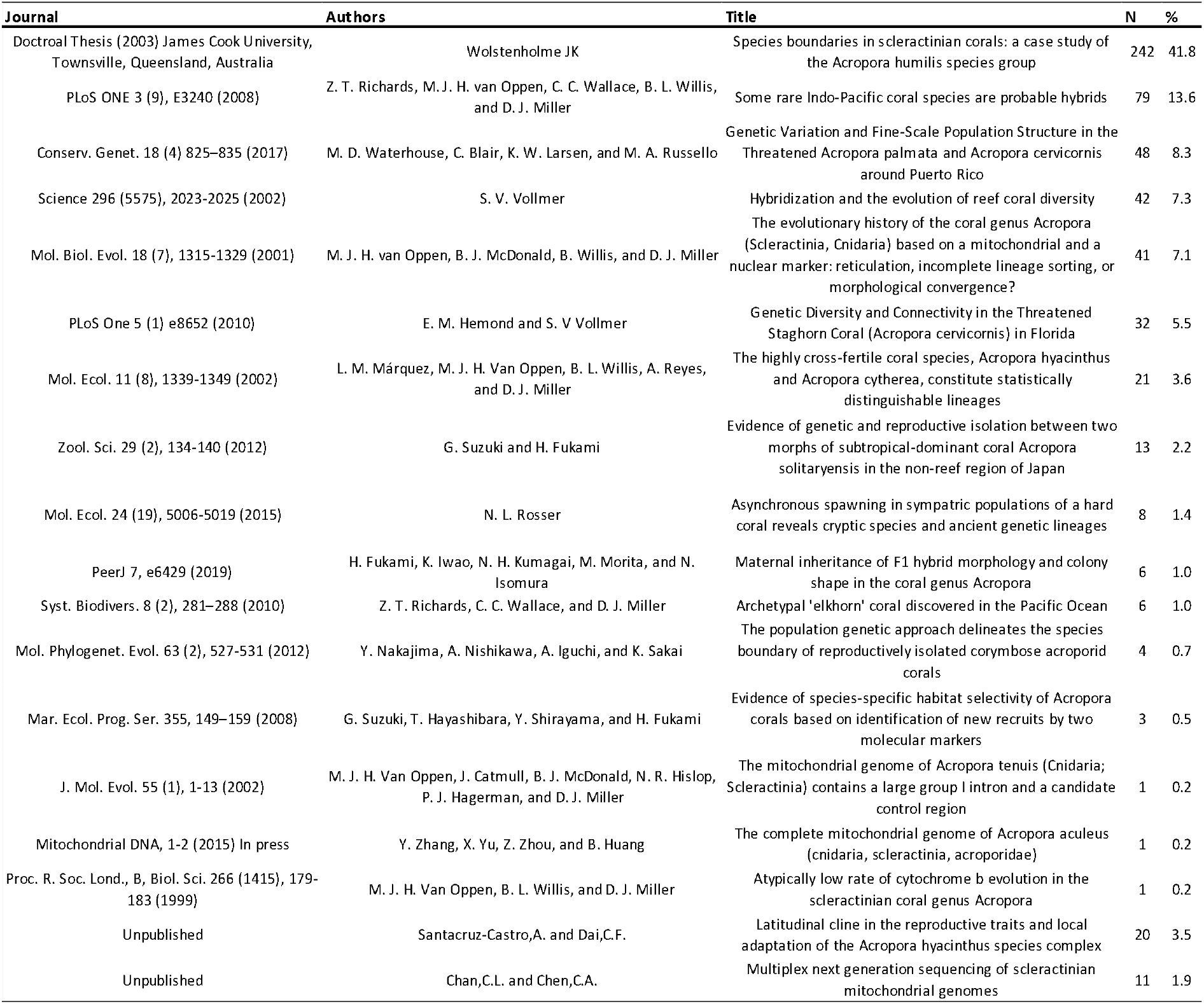
Sources of mtCR references sequences by Paper title

**Table 4:**
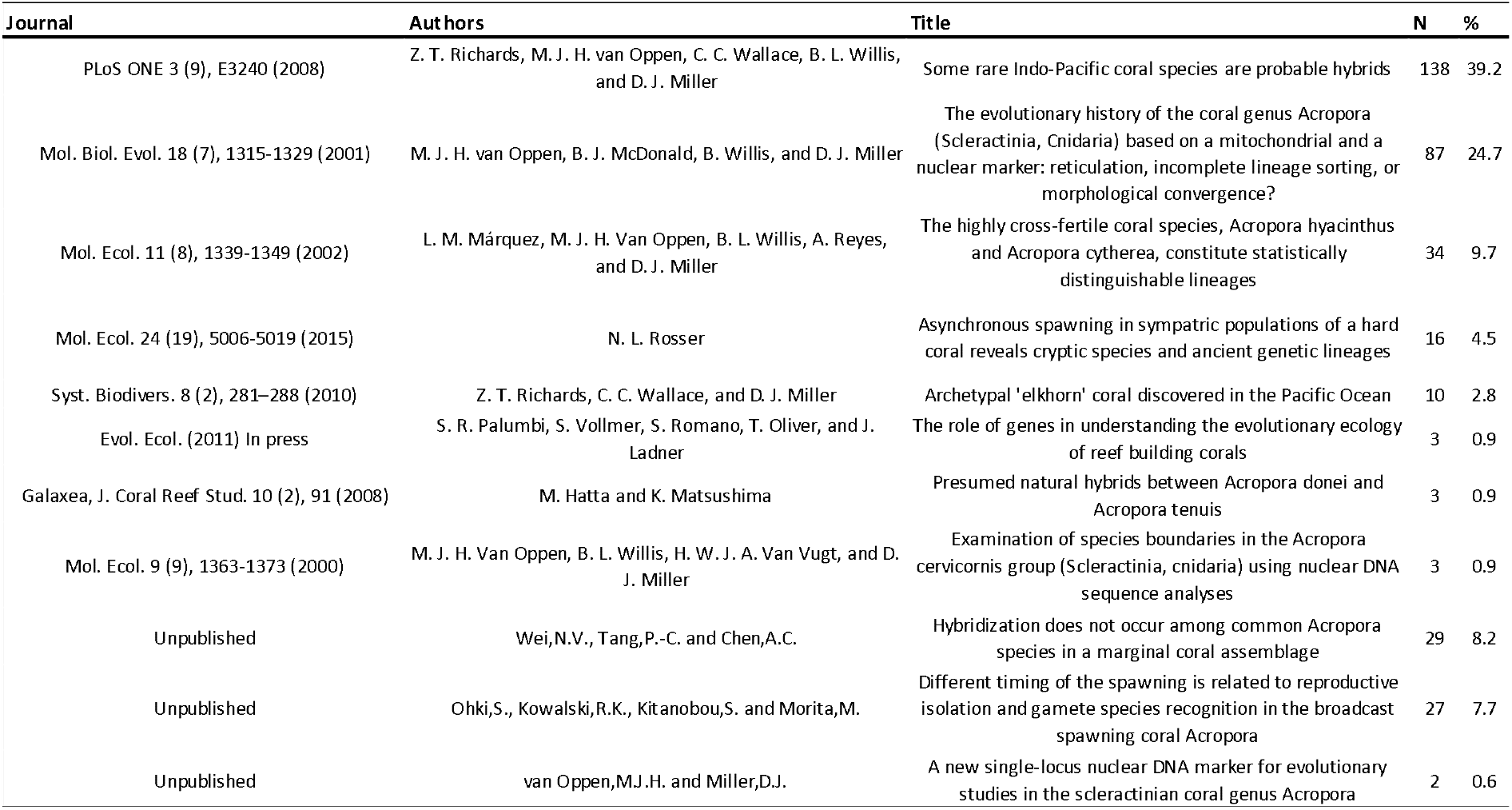
Sources of PaxC references sequences by Paper title

### Phylogeny

While the construction of phylogenetic trees is a more refined and, by some, recommended form of species identification, it is more demanding in terms of time and expertise, and has stages such as sequence alignment and phylogenetic interpretation that often require significant manual interpretation and intervention (Nixon and Wheeler 1990; Minh et al. 2013; Nguyen et al. 2015). We observed high levels of polyphyly across all of our highest confidence matches, with only 1 of evident case in which the 100% match was to a monophyletic species (*Acropora humilis* PaxC region). Our preference is to employ a standardised, quickly repeatable, objective process, while keeping in mind that the results are imperfect and designed to inform but not dictate identification efforts.

### Blast match

The use of query match with blast is far from being a complete solution, with 26 out of 76 samples without a single species or species group resolved ID (Table 5), we have proposed a theoretical decision tree that standardise the choice when there is a need of sorting through the output of a multi-species name blast match (Fig. 2). In addition we used two different regions, a mitochondrial one and a nuclear one to increase the confidence in our results (Suzuki et al. 2008). Our proposed species ID are provisional and subject to revision when more species and sequences are available, hence the value of a repeatable process for matching and re-matching.

All methods of identification that rely on DNA barcodes (e.g.: Blast matching, Barcoding Gap analysis, etc.) use genetic divergence thresholds to assign individuals to correct species. This based on two fundamental assumptions 1) monophyly of species with respect to the molecular marker used, and 2) intraspecific genetic divergence is much smaller than interspecific genetic differences (Toffoli et al. 2008). The first assumption is more dependent on the experimental choices made (i.e. sequence region selected). The second assumption is where most of the criticism of barcodes methods lies, on not comparing sister species or geographical distribution thus, underestimating intraspecific genetic variability (Moritz and Cicero 2004; Toffoli et al. 2008). Fundamentally, the issue of using thresholds lies in the fact that species naturally embodies an evolutionary process, being subject to demographic and selective processes that will act on the genetic diversity (e.g. (Avise et al. 2000; Hey 2001; Coyne and Orr 2004; Toffoli et al. 2008)). Since species are real evolutionary groups and not categories which are created as a direct function of perceived distinction (Hey 2001), the use of thresholds in “discovering” new species is overly simplistic, and in some cases even misleading (Toffoli et al. 2008). We argue against simply substituting morphology based taxonomy with DNA barcoding based taxonomy, as each system is more adapt to answer different questions (Toffoli et al. 2008). Correct assignment is thus only possible by complementing DNA barcoding with other data types, such as morphological, and ecological characters as part of an integrated diagnostic approach (Toffoli et al. 2008).

While the simple blast match is inheritably less complex than other traditional DNA barcodes identification methods (i.e.: the above mentioned barcoding gap analysis and, the potentially more direct Automatic Barcoding Gap Discovery (ABGD) (Puillandre et al. 2012)). The argument we make is that given the shared limitation and the common need for additional data in an integrated approach, a faster and simpler blast match provides a relatively similar additional amount of information for the species identification while being more easily repeatable, especially considering the value of standardized data among aquariums (da Silva et al. 2019) and in light of the ever increasing quantity of reference materials being added to GenBank and BOLD.

Available sequences libraries are not exhaustive, and probably never will be. That should not stop us from attempting identifications based on the best available knowledge. All identifications must fit within the context of our evolving understanding of taxonomy, where new descriptions invalidate existing ones and often we do not have unambiguous taxonomic descriptions for every species (Simpson 1951; Wiley 1978; Mayr 1982, 2000; Nixon and Wheeler 1990; Medlin et al. 1995). The reality is that barcoding as an independent tool of identification has its clear limitation (Van Oppen et al. 1999; Shearer et al. 2002; Ferguson 2002; Moritz and Cicero 2004; Lee 2004; DeSalle et al. 2005; Meyer and Paulay 2005; Prendini 2005; Knapp et al. 2005; Cognato 2006; Hickerson et al. 2006; Meier et al. 2006, 2008; Little and Stevenson 2007; Kitahara et al. 2010), and coral barcoding is no exception. With some of the concern on the use of genetic repertories in common of all the methods explored in this study, we strongly recommend an integrated diagnostic approach, combining morphological and genetic means of identification, in agreement with C. Moritz in “DNA Barcoding: Promise and Pitfalls” (Moritz and Cicero 2004). Proposing the use of barcoding and query matching as an additional tool for identification, increasing confidence of experts and both confidence and capacity of non-experts’ taxonomist. Producing this way reliably identified sequences that can potentially become new reference, adding to the repertories already freely available. Considering the specific needs of the aquarium community and their critical role to support corals conservation (i.e., by hosting at least 24% of the corals assessed as highly vulnerable to climate change (da Silva et al. 2019)), we emphasise the value of integrating standardised barcoding analysis, and specimen morphology photographs to optimise usage of authoritative identification guides and expert opinion.

## Acknowledgments

Genetic analysis costs were partly covered by a FAITAG genetics fund grant an initiative from the EAZA (European Association of Zoos and Aquaria) and EUAC (European Union of Aquarium Curators) Fish and Aquatic Invertebrate Taxon Advisory Group (FAITAG).

